# Roads shape seed dispersal by rodents and carnivores at forest edges

**DOI:** 10.1101/2025.03.19.644142

**Authors:** João Craveiro, Miguel N. Bugalho, Pedro G. Vaz

## Abstract

The influence of road networks on seed dispersal remains poorly understood. We conducted a field experiment in Mediterranean oak woodlands of southern Portugal to assess how road type (paved vs. unpaved) and road-forest context (edge vs. non-edge) shape seed dispersal by rodents and carnivores. Using labeled acorns and seed mimics, we tracked dispersal distances, number of dispersals, road crossings, and dispersal directions. Rodents dispersed seeds farther in forest edges and along paved roadsides, predominantly moving them parallel to roads, but rarely across—although crossings were more frequent on unpaved roads. In contrast, carnivores mediated long-distance dispersal, primarily perpendicular to roads, and although seed road crossings were rare, carnivore-dispersed seeds crossed roads—especially unpaved ones—nine times more frequently than rodent-dispersed seeds. Shrub cover increased rodent-mediated dispersal events, heavier acorns traveled farther, and water patches decreased carnivore-mediated dispersal events near roads. These findings highlight the dual role of roads as barriers and of roadsides as corridors for seed dispersal, with implications for forest regeneration and landscape connectivity. Roadside management should prioritize moderate shrub cover along paved roads to support rodent dispersal while balancing ecological benefits with potential trade-offs. Unpaved roads, in turn, should be managed as functional corridors for carnivores, enhancing connectivity between fragmented forests. By integrating species-specific dispersal processes into road planning, landscapes can be designed to minimize road barrier effects while promoting natural regeneration in human-modified forests.

## 1. INTRODUCTION

Seed dispersal is a vital ecological process shaping plant population dynamics and recruitment (Nathan & Muller-Landau, 2000; Wang & Smith, 2002; Chen et al., 2019b). Terrestrial mammals, particularly rodents and carnivores, drive seed dispersal through scatter-hoarding (storing seeds in small, dispersed caches; Gómez et al., 2019; Vaz et al., 2024) and endozoochory (ingestion and deposition; Salgueiro et al., 2019; Rubalcava-Castillo et al., 2020). While rodents typically disperse seeds over short distances, carnivores transport them farther due to their larger home ranges (Herrera et al., 2015). However, human-induced landscape changes, especially roads, may disrupt these plant-animal interactions by altering animal movement and landscape connectivity (Markl et al., 2012; Ibisch et al., 2016; Cui et al., 2018; Quiles & Barrientos, 2024).

The position of a road relative to a forest—whether crossing its interior or delimiting its edge—can further affect seed dispersal patterns. These road-forest contexts influence mammal behavior, seed dispersal rates, and movement patterns, with potential cascading effects on plant regeneration and ecosystem resilience (López-Barrera et al., 2007; Mazzamuto et al., 2018). Yet, their role in animal-dispersed forest systems remains largely overlooked, including in Iberian evergreen oak woodlands, where severe oak decline and lack of natural regeneration has been reported since the early 1990s (Bugalho et al., 2011; Vaz et al., 2019). These oak systems provide key ecosystem goods, such as fuelwood, game, crops, and cork (Bugalho et al., 2009), alongside biodiversity conservation benefits (Bugalho et al., 2011; Plieninger et al., 2011), but face threats from climate change (Díaz et al., 2021), pathogens (Branco & Ramos, 2009; Brasier, 1992, 1996), phytophagous insects (Branco et al., 2002), wildfires (Moreira et al., 2011; Vaz et al., 2013), and overgrazing (Wadud et al., 2024). Understanding seed-dispersal dynamics by mammals such as rodents and carnivores, is crucial for promoting adequate oak natural regeneration in these ecosystems.

Rodents are essential acorn dispersers, collecting and storing acorns in various microhabitats (Evrard et al., 2017; Vaz et al., 2024). While many are later retrieved and consumed, others remain buried, germinating and contributing to forest regeneration (Perea et al., 2011a). Rodents also frequently use roadside verges as refuges, especially where the surrounding landscape is managed for livestock grazing or wildfire prevention (Ruiz-Capillas et al., 2013; Galantinho et al., 2020). Carnivores, in turn, are major dispersers of fleshy-fruited species, benefiting wildlife, livestock, and ecosystems with human value (Pinto-Correia et al., 2011; Moreno et al., 2018; Rubalcava-Castillo et al., 2021). They transport seeds over long distances, often dispersing them along roadsides used as movement corridors (Suárez-Esteban et al., 2013a).

Roads can create barriers to animal movement or alter seed dispersal behavior (Suárez-Esteban, et al. 2014). Some studies suggest that roads reduce rodent-mediated dispersal distances (Chen et al., 2019b), while differences in road tolerance among mammals can lead to changes in activity patterns and resource use (Quiles & Barrientos, 2024). Research in southern Spain suggests that unpaved roads promote carnivore-mediated seed dispersal along road verges (Suárez-Esteban et al., 2013a), whereas in China, paved roads significantly influence rodent-mediated seed dispersal distance and removal rates, with differences between forest interiors and roadsides (Niu et al., 2018, 2021). Notably, seed road crossings by terrestrial mammals remain largely undocumented.

Edges, as transitional zones between habitats, further shape seed dispersal by mammals. López-Barrera et al. (2007), in the highlands of Chiapa, in Mexico, found reduced acorn removal in grasslands with pronounced edges, while more dispersal events occurred from forest edges into interiors. Similarly, Herrera et al. (2015), in an oak woodland in Alentejo, Southern Portugal, showed that amount and spatial configuration of tree cover affect carnivore-mediated seed dispersal distances. These results suggest that habitat shifts, such as those at edges, may determine seed dispersal depending on species or functional group.

Despite their potential impact, roads’ effects on mammal-mediated seed dispersal remain poorly understood. We conducted a field experiment in an oak woodland area in southern Portugal to assess how road-forest context (edge vs. non-edge) and road type (paved vs. unpaved) may influence seed dispersal. We examined dispersal distance, number of dispersals, seed road crossings, and dispersal directions relative to roads, mediated by rodents and carnivores. We asked:

i. How do dispersal distance and the number of dispersals vary with road-forest context and road type in rodents and carnivores? We hypothesized that rodents would disperse seeds more often and over longer distances in edge than non-edge contexts, where acorn availability is lower. For carnivores, we expected greater dispersal distances and more dispersal events in non-edge contexts, reflecting their preference for forested areas. As unpaved roads pose less of a barrier, we also hypothesized that both groups would disperse seeds more frequently and over longer distances along unpaved than paved roads.
ii. What proportion of dispersed seeds cross the road in rodent- and carnivore-mediated dispersals? We hypothesized that carnivores, with their larger home ranges and greater mobility, would mediate more seed road crossings than rodents.
iii. Do seed dispersal direction relative to roads differ between rodents and carnivores? We hypothesized that rodents would disperse seeds mainly along roadsides, as they use road verges as refuges and have small home ranges, whereas carnivores would disperse seeds away from the road due to their larger home ranges and broader movements.

## 2. METHODS

### 2.1 Study area and main study design

We conducted the study on 12 road sections (1.2 km each) in the Alentejo region, southern Portugal (Fig. 1). The local climate is Mediterranean, with cold, wet winters and hot, dry summers (mean annual precipitation: 620 mm; mean temperature: 18 °C). The landscape consists of undulating terrain (200–400 m asl) dominated by *montado* (*dehesa* in Spain) and pastures. Montados are characterized by an open-canopy of cork (*Quercus suber*) and holm (*Quercus rotundifolia*) oak stands with an understory of shrubs (e.g., *Cistus*, *Ulex*, *Erica*, *Lavandula* spp.; Bugalho et al., 2009, 2011; Castro & Freitas, 2009; Azedo et al., 2022) and grassland areas. Grasslands in open areas consist mainly of annual forbs, graminoids, and legumes. Livestock, mainly cattle and sheep, with occasional domestic pigs, occurs in the region. Acorns naturally fall between September and January.

**Figure 1.**
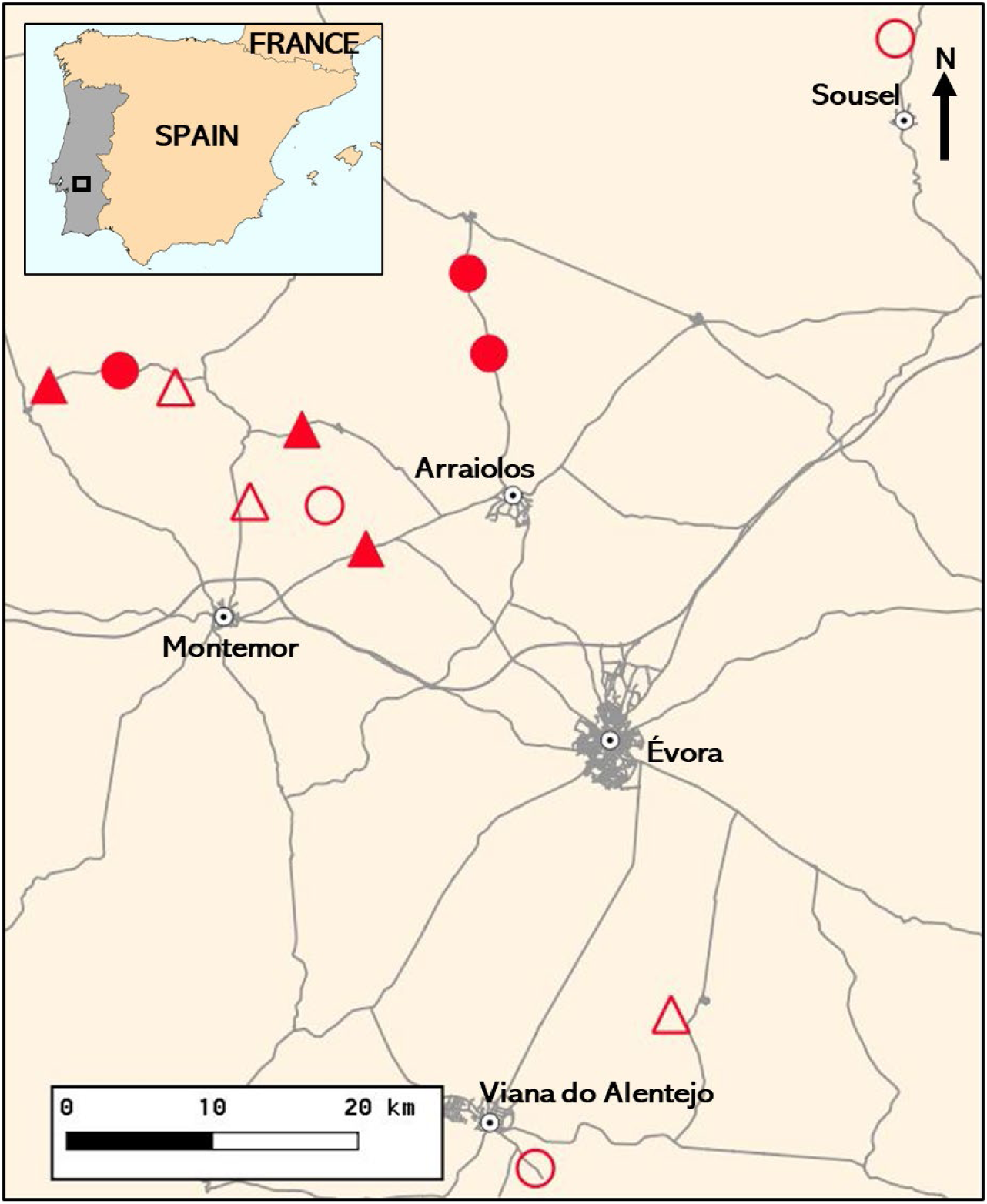
Location of the 12 road sections (1.2 km each) in the Alentejo region, southern Portugal, where seed dispersal by rodents and carnivores was monitored. The experimental design included two treatments: road-forest context (edge [circles] vs. non-edge [triangles]) and road type (paved [solid symbols] vs. unpaved [unfilled symbols]).

The 12 road sections (≥1.5 km apart) encompassed two road-forest contexts—six bordering oak forests (edge) and six traversing oak forests (non-edge)—and two road types (six paved, six unpaved). Edge roads were characterized by oak woodland on one side and open pasture or cropland on the other. Paved roads (∼8 m wide; range: 6–11 m) were National Roads with asphalt pavement and a traffic volume of 95 vehicles/hour (range: 19–234) (see www.infraestruturasdeportugal.pt). Unpaved roads (∼4 m wide; range: 3–7 m) were public dirt roads with low traffic, mainly for agricultural access. To account for spatial variation, we established three replicates per factor (2 road-forest contexts × 2 road types × 3 replicates = 12 road sections) (Fig. 1, Fig. 2).

**Figure 2.**
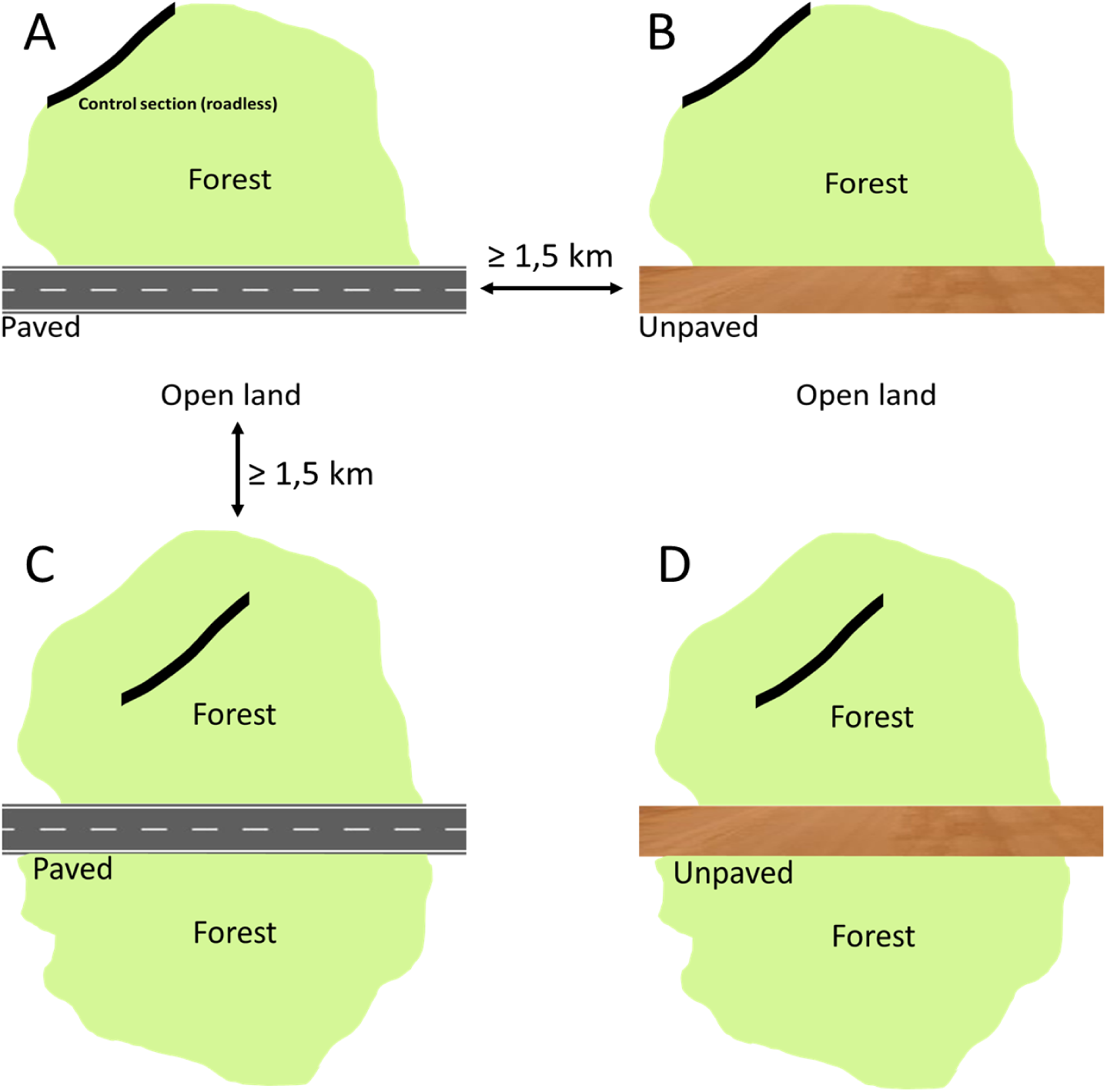
Schematic representation of the road-forest contexts and road types. (A, B) Edge sites with different road types, where the forest borders a tree-free open area, typically used for pasture or cropland. (C, D) Non-edge sites with different road types traversing a continuous forest. Solid lines indicate the corresponding roadless sections: in edge sites, these sections bordered the oak forest, whereas in non-edge sites, they extended through the forest.

The main acorn-dispersing rodents in the area are the wood mouse (*Apodemus sylvaticus*) and the Algerian mouse (*Mus spretus*) (Galantinho et al., 2020). Several carnivore species in the study area contribute to fleshy-fruit dispersal, including stone marten (*Martes foina*), European badger (*Meles meles*), red fox (*Vulpes vulpes*), Egyptian mongoose (*Herpestes ichneumon*), common genet (*Genetta genetta*), and Eurasian otter (*Lutra lutra*) (Craveiro et al., 2019).

### 2.2 Assessing rodent-mediated seed dispersal

To control for road effects on rodent-mediated seed dispersal, each of the 12 sites included a comparable roadless section. In edge sites, this section bordered the oak forest, while in non-edge sites, it traversed the forest (Fig. 2).

To evaluate the effects of road-forest context and road type on rodent-mediated seed dispersal, we tracked 1440 labeled holm oak acorns from November 2022 to May 2023. One week before tracking, ∼3000 ripe acorns were collected from six holm oaks per site, either from the ground or directly from trees if they detached easily. Acorns were selected based on uniform brown color and absence of desiccation, then mixed and floated in water to remove weevil-infested ones. Mean pre-labeling acorn weight was 5.44 g (± 0.04 SE, range = 2.10–16.27 g). Acorns were labeled by drilling a 1-mm hole (avoiding the embryo) and inserting a 3-cm wire (0.5 mm diameter) attached to a 14.5 × 1.6 cm yellow numbered plastic label (Chen et al., 2019a). Labeling added 6–20% to acorn weight. This method, which does not alter dispersal patterns, ensures high recovery rates (Xiao et al., 2006).

Each road and roadless section had six acorn supply stations—three per side of the road (or imaginary road in roadless sections)—spaced ∼200 m apart. Along roads, stations were placed 3–10 m from the roadside (asphalt for paved roads, dirt for unpaved). Roadside vegetation consisted of narrow strips of herbaceous plants, shrubs, and occasionally isolated trees, mostly oaks. Across the 12 sites, this resulted in 72 supply stations along roads and 72 in roadless sections. To exclude non-rodent consumers, each station consisted of a 32 × 25 × 12 cm wire cage (1.4 cm mesh), containing 10 labeled acorns. Each cage had two 5 × 5 cm entrances and was secured to prevent displacement (Perea et al., 2011a, Vaz et al., 2024). Stations were placed within 40 cm of a shrub, mainly *Cistus* spp or *Ulex* spp (90% of cases). To standardize accessibility, vegetation inside cages was cleared, leaving acorns exposed.

Acorn dispersal was tracked within a minimum 50 m radius around supply stations, expanding by 50 m when a dispersed acorn was found (Vaz et al., 2024). Monitoring followed a decreasing frequency: every 2 days up to day 8, then twice a week for two weeks, weekly for two weeks, biweekly for one month, monthly for two months, and finally bimonthly for four months. Each acorn’s new location was recorded. Depredated acorns (embryo damaged) and those missing (i.e., not found in subsequent searches) were assigned their last recorded dispersal distance. Acorns predated inside supply stations were excluded from dispersal analyses. From week one, non-viable acorns (shriveled, yellowed, hollow) were also removed (Vaz et al., 2024). Some cattle and sheep were present at three sites during the final three weeks of the experiment near roadless sections, not road verges, but had minimal impact on seed tracking.

Dispersal distance (from supply station) and distance from the road were measured using a laser meter (1-mm precision) or tape measure. To assess the effect of acorn arrival microhabitat (Perea et al., 2011b; González-Rodríguez & Villar, 2012; Vaz et al., 2024), deposition sites were classified as open areas or under shrubs. Within a 50 m radius of each station, on the same side of the road, we quantified tree density (trees/ha) by counting trees and visually estimated shrub and grass cover (%) (Galantinho et al., 2020).

To estimate natural acorn availability, we counted acorns beneath the three nearest oak trees to each station (Vaz et al., 2024). Counts were conducted at four cardinal points under the canopy, facing the trunk, with 30 seconds per point. The sum per tree was averaged across the three trees for a station-level estimate. A single observer conducted all counts in early 2023, after acorn drop.

To examine rodent density effects on acorn dispersal, we conducted a 4-night live-trapping session at acorn supply stations (Perea et al., 2011c Niu et al., 2021). Four Sherman traps were placed in favorable microsites to maximize captures and improve abundance estimates (under shrubs *Cistus spp*, *Ulex* spp or tall grass; Fedriani, 2005) at increasing distances from the station (0 m, 10–15 m, 20–30 m, 40–50 m). Traps were baited with peanut butter on toast (4 × 4 cm), and cotton wool was provided for insulation. Captured rodents were marked with a small lateral fur cut (Hoffmann et al., 2010; Jumeau et al., 2017) and released at capture sites. Procedures complied with Directive 2010/63/EU on animal use in research.

### 2.3 Assessing carnivore-mediated seed dispersal

Due to their large home ranges and the widespread presence of roads, carnivore-mediated seed dispersal could not be sampled in roadless control sections as for rodents. To assess the influence of road-forest context and road type, we supplied dried common figs (*Ficus carica*), previously used in feeding experiments (Herrera et al., 2015). The figs were distributed across the 12 road sections from October 4, 2022 (fall) to July 18, 2023 (summer), ensuring proportional representation of each road-forest context and road type to avoid seasonal biases.

Each road section had six supply stations per side, ∼200 m apart along the same side, totaling 144 stations. At each station, we placed six aluminum trays (14.5 × 12.0 × 4.0 cm) spaced 10 m apart, each containing six dried figs. Every fig embedded 10 plastic beads uniquely color-coded for each station (seed mimics, 2.6 mm in diameter) (Herrera et al., 2015; Salgueiro et al., 2019). To document carnivore consumption of figs, we placed two Bushnell CORE S-4K model 119949C cameras at two randomly selected stations.

After eight days, we systematically searched for carnivore scats containing seed mimics within a 1200 m buffer around the supply stations, corresponding to the average home range radius of the carnivore community (Dekker et al., 2001; Planillo et al., 2018; Craveiro et al., 2019). The search focused on carnivore-utilized paths (e.g., trails, firebreaks, animal tracks; Herrera et al., 2015), covering on average 88% of the paths per buffer (range: 81%–98%). Scat locations were geo-referenced using a Garmin Dakota 10 GPS (mean error ± 3 m). Dispersal distance was calculated in GIS as the straight-line distance between the supply station and the scat location. Seed mimics from each scat were matched to their supply station of origin using their unique color code. We also calculated the distance from the scat location to the road and determined whether a ‘seed’ road crossing had occurred.

Tree density (oak trees/ha) was estimated within each 1200 m buffer around road sections, as this may influence carnivore habitat use. Using GIS, we generated five random points per road side and created 50-m radius buffers around each, totaling 10 buffers per site. The number of oaks per buffer was counted via satellite imagery, and density was calculated by dividing the total number of trees by the combined buffer area. This random sampling approach avoided the need for full tree counts within the 1200 m buffer.

To account for variations in carnivore activity within the buffer, we calculated the Kilometer Abundance Index based on all carnivore scats found along the walked paths, including those with and without seed mimics (Vincent et al., 1991). This index served as a proxy for carnivore abundance to help interpret seed dispersal patterns.

Additionally, we quantified landscape characteristics within the 1200 m buffer using QGIS (v3.20.3-Odense), considering total stream length and the number and total area of water patches—variables selected for their potential relevance to carnivore-mediated seed dispersal, as indicated by preliminary analyses.

### 2.4 Analyses

#### 2.4.1 Rodent-mediated seed dispersal – distance and number of dispersals

To assess the effects of road type, road-forest context, and covariates (Table 1) on acorn dispersal distance and number of dispersals (i.e., acorns moved per supply station), we used generalized linear mixed models (GLMMs) with station and site as random factors. Correlation matrices showed no collinearity issues among predictors (*r_s_* ≤ 0.58).

**Table 1.** Names, definitions, and range of variables considered in the analysis of rodent- and carnivore-mediated seed dispersal distance and number of dispersals.

| Group | Variable | Description | Range |
| --- | --- | --- | --- |
| Rodents | Road-forest context | Position of the road relative to the forest: "edge" (oak forest on one side, open area on the other) or "non-edge" (road traversing oak forest) |  |
|  | Road type | Road surface type: "paved" (asphalt) or "unpaved" (dirt road) |  |
|  | Road effect | Road sections vs. adjacent roadless control areas with the same forest context (edge or non-edge); categorized as "road" vs. "roadless". |  |
|  | Distance from road | Shortest distance (m) from the acorn's final location to the road. |  |
|  | Acorn weight | Weight (g) of each acorn. | 2.10–16.27 |
|  | Rodent density | Estimated (individuals/ha) separately for road and roadless sections within each site. | 0.95–7.45 |
|  | Acorn availability | Mean number of acorns per station, estimated by visual counts at each of the three nearest oaks. | 0.00–224.67 |
|  | Arrival microhabitat | Deposition microsite of each acorn (inside a shrub vs. open area). |  |
|  | Shrub cover | Visually estimated shrub cover (%) within a 50 m radius of each supply station. | 0–97 |
|  | Grass cover | Visually estimated grass cover (%) within a 50 m radius of each supply station. | 1–100 |
|  | Tree density | Tree density within a 50 m radius of each supply station. | 0.00–227.50 |
| Carnivores | Road-forest context | See above. |  |
|  | Road type | See above. |  |
|  | Distance from road | Shortest distance (m) from the seed's final location within a carnivore scat to the road. |  |
|  | Carnivore abundance | Kilometric abundance index of carnivore scats | 0.69–3.01 |
|  | Length of streams | Total length of watercourses (m) within a 1200 m buffer. | 4103–9597 |
|  | Number of water patches | Number of water patches. | 12–32 |
|  | Area of water patches | Total area (ha) occupied by water patches within. | 1.68–20.74 |
|  | Tree density | Number of oaks per hectare estimated from ten 50-m radius buffers (five per road side). | 17.60–55.40 |

Rodent density was estimated separately for road and roadless sections using ‘Rcapture’ (v1.4-4; Baillargeon & Rivest, 2009), assuming closed populations (i.e., no mortality or migration during the sampling period). We applied the “closed.bc” function to refine estimates and correct bias in capture–mark–recapture data.

For dispersal distance, we used a GLMM with a zero-inflated Gamma distribution and log link function, with each acorn’s final position as the response variable. For number of dispersals, we fitted a GLMM with a Conway–Maxwell-Poisson (compois) distribution and log link, which accounts for overdispersion or underdispersion in count data. Although each station had a single observation, including station as a random effect (site/station) improved the number of dispersals model fit (Shmueli et al., 2005; Lynch et al., 2014).

The final models were selected through a three-step process. First, we fitted univariate GLMMs for each predictor and selected for inclusion only those that significantly improved model fit (*P* < 0.05) compared to the null model using likelihood ratio tests (LRTs; e.g., Azedo et al., 2022). Road-forest context, road type, and road effect were included a priori due to their importance to the hypotheses. Second, full models were fitted with selected predictors. Fixed effects for dispersal distance included road-forest context, road type, their interaction, road effect, arrival microhabitat (shrub/open), distance from road, acorn weight, and shrub cover. For number of dispersals, arrival microhabitat was excluded, and rodent density was included. Finally, we used backward elimination, sequentially removing non-significant predictors based on LRTs for nested models (Zuur et al., 2009).

#### 2.4.2 Carnivore-mediated seed dispersal – distance and number of dispersals

To assess predictors of ‘seed’ dispersal distance, we used a GLMM with a Tweedie distribution and log link function, modeling the straight-line distance between each supply station and the scat containing seed mimics as the response variable. The Tweedie distribution accommodates non-negative continuous data and supports a wide range of distributions, including gamma and Poisson-gamma mixtures. Site was included as a random factor. Correlation matrices indicated no collinearity issues (*r_s_* ≤ 0.68).

To examine predictors of the number of dispersals, we fitted a GLMM with the count of ‘seeds’ dispersed per site as the response variable. Given the over-dispersed nature of the data, we applied a lognormal Poisson family distribution with a log link, including site as a random effect (e.g., Garcia et al., 2022).

Final models were selected through a three-step procedure, as described for rodents. First, we fitted univariate GLMMs and retained only predictors that significantly improved model fit compared to the null model (LRTs). Road-forest context and road type were included a priori. Second, full models were constructed with selected predictors. Fixed effects for dispersal distance included road-forest context, road type, their interaction, distance from road, number of water patches, and area of water patches. Carnivore abundance, total length of streams, and tree density were excluded, as their univariate models did not improve the null model fit. For the number of dispersals, we included the same categorical variables, plus total length of streams and number of water patches, excluding carnivore abundance, area of water patches, and tree density for the same reason. Third, backward elimination was applied, sequentially removing non-significant predictors using LRTs for nested models.

All models were implemented in the ‘glmmTMB’ R package (v1.1.7; Brooks et al., 2017). Model adequacy was assessed using the ‘DHARMa’ R package (v0.4.6; Hartig, 2022), which generates residual diagnostics, including QQ-plots, dispersion tests, and residuals vs. predicted values plots. Analyses were conducted in R version 4.1.1 (R Core Team, 2021).

#### 2.4.3 Seed dispersal directions by rodents and carnivores relative to the road

To assess whether seed dispersal directions differed between rodents and carnivores, we determined the angular deviation of each dispersal event from the road. Angles were measured as the deviation between the straight-line dispersal trajectory (from the supply station to the deposition site) and the road axis, with 0° indicating movement parallel to the road and 90° indicating movement perpendicular to it. As the number of dispersed acorns exceeded that of fleshy-fruit ’seeds,’ we analyzed the relative proportions of dispersal events to ensure comparability. A wind rose plot was used to visualize directional trends.

## 3. RESULTS

Seed dispersal patterns varied markedly between rodents and carnivores in terms of dispersal distance, number of dispersals, road crossings, and dispersal directions. Both road-forest context and road type influenced some of these differences.

### 3.1 Rodent-mediated seed dispersal

Rodents dispersed 744 of 1,440 acorns (52%), with a mean dispersal distance of 4.7 m. Dispersal was consistently greater near road sections than in roadless controls, regardless of road-forest context or road type (Table 2). Only 22 acorns (3%) crossed roads, a proportion comparable to roadless controls (24 acorns). Crossings were evenly distributed between edge and non-edge contexts (11 each) and were 2.1 times more frequent on unpaved (15) than paved roads (7). Unpaved roads had three times more crossings than their controls (15 vs. 5), whereas paved roads had only 37% of the crossings observed in controls (7 vs. 19).

**Table 2.**
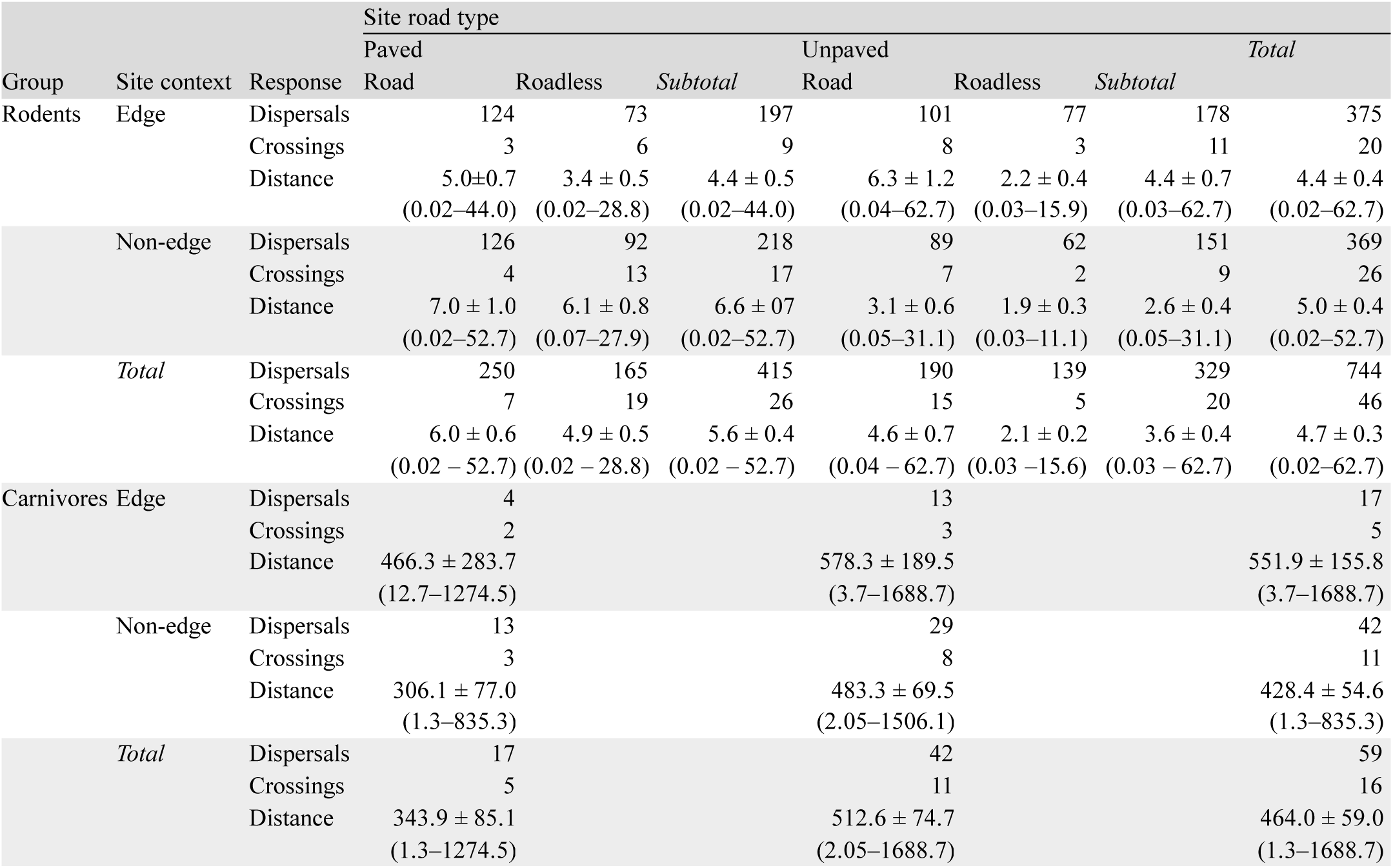
Observed seed dispersal metrics for rodents and carnivores across edge and non-edge road-forest contexts with paved and unpaved roads. Shown are the total number of dispersals, seed road crossings, and dispersal distances (m) (mean ± SE, range). For rodents, values are provided for road sections and their paired roadless controls, whereas no roadless control was available for carnivores.

Wood mice (80% of captures) and Algerian mice (20%) mediated most dispersals, with a combined estimated density of ∼3.4 individuals ha⁻¹. Rodent density was higher near roads (1.6 times) than in roadless controls. By the end of the experiment, 87% of dispersed acorns had been consumed, only 3 (0.4%) had emerged as seedlings, and 26% of the 1,440 offered acorns had gone missing.

#### 3.1.1 Rodent-mediated seed dispersal distances

Accounting for random effects (supply station and site), our optimal mixed model (Table 3) confirmed that both road-forest context and road type significantly influenced dispersal distances (Fig. 3). As hypothesized, acorns dispersed 1.7 times farther in edge than in non-edge contexts. However, contrary to expectations, dispersal distances at unpaved road sites were 42% shorter than at paved road sites. Within sites, dispersal distances were 1.6 times greater near road sections than in roadless controls.

**Figure 3.**
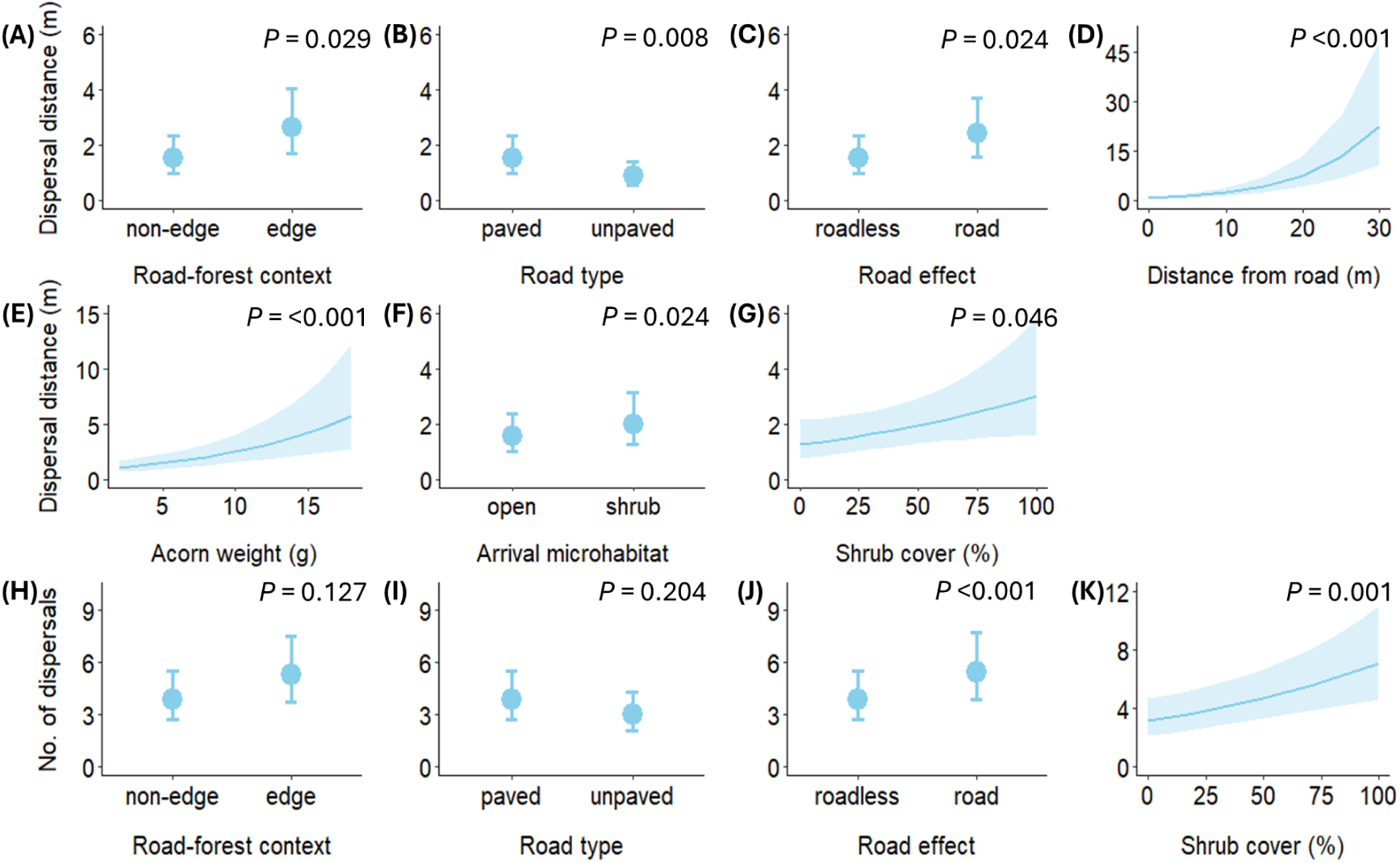
Mean fitted values (±95% CI) from the optimal mixed-effects models predicting (A–G) acorn dispersal distance and (H–K) number of acorn dispersals mediated by rodents.

**Table 3.**
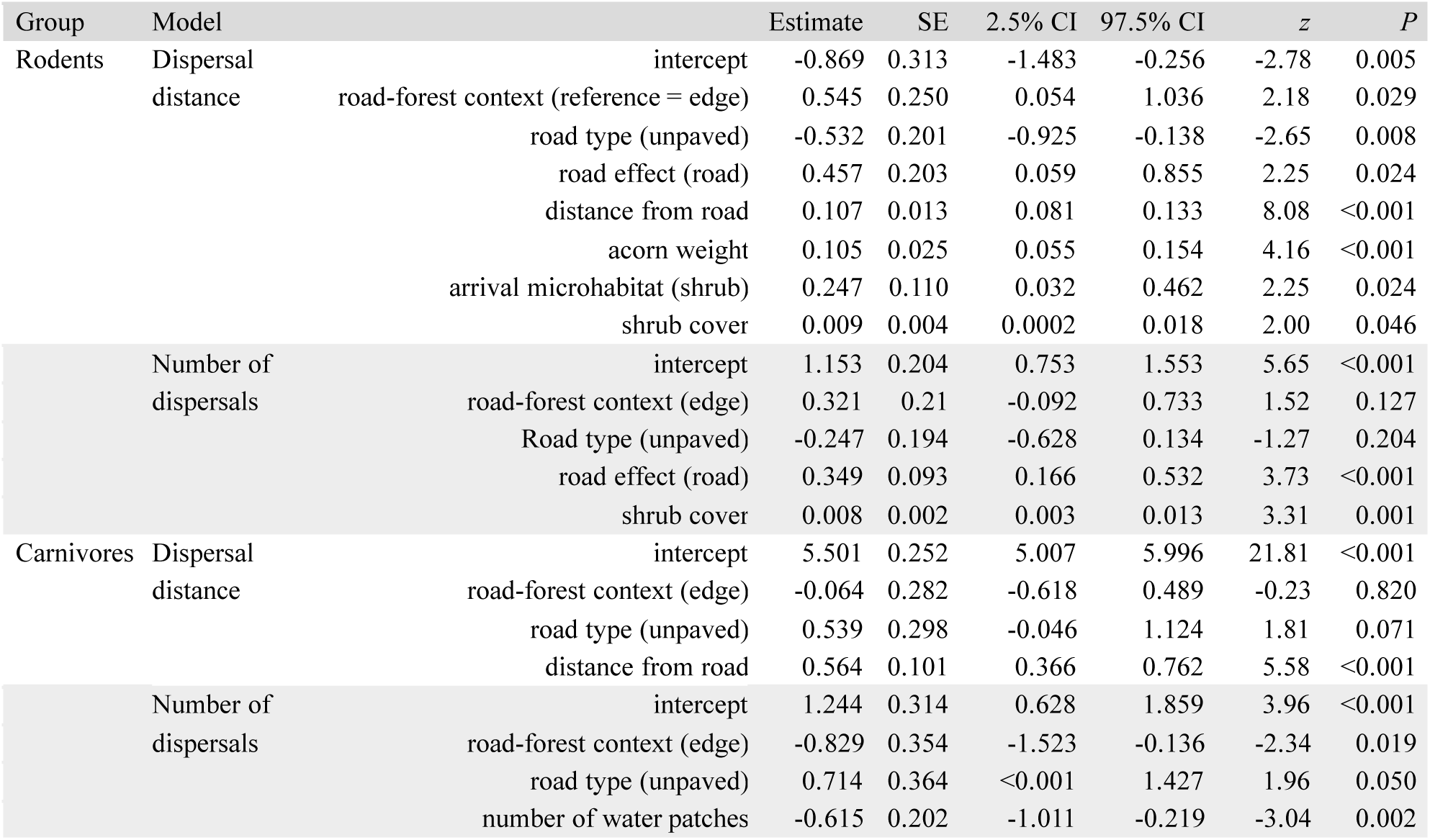
Parameter estimates (Estimate ± SE), 95% confidence intervals (CI), *z*-values, and *P*-values for the fixed effects in the final mixed-effects models predicting acorn dispersal distance and number of dispersals by rodents.

Dispersal distance also increased with distance from the road: acorns recovered 20 m from the road traveled 5 times farther than those at 5 m. Heavier acorns traveled farther, with a 68% increase in dispersal distance for acorns weighing 10 g compared to 5 g. Shrub cover further enhanced dispersal: acorns deposited under shrubs traveled 28% farther than those in open areas, and increasing cover from 25 to 50% resulted in a 24% rise in dispersal distance.

#### 3.1.2 Number of rodent-mediated seed dispersals

When accounting for random effects, neither road-forest context nor road type significantly influenced the number of acorns dispersed, providing no evidence to support our hypotheses (Table 3; Fig. 3). However, road presence significantly increased dispersal, with 1.4 times more acorns dispersed near road sections than in roadless controls. Additionally, shrub cover had a positive effect: increasing cover from 25 to 50% led to a 22% rise in the number of acorns dispersed.

### 3.2 Carnivore-mediated seed dispersal

Across ∼350 km of systematic scat surveys at the 12 sites, we recorded 612 carnivore scats, of which 41 contained fig ‘seeds’, corresponding to 59 dispersal events (Table 2). The mean dispersal distance from the supply station was 464 m.

As predicted, carnivores mediated proportionally more seed road crossings than rodents. Of the dispersed ‘seeds’, 27% (16) crossed roads—nine times the percentage observed for rodents (3%). Crossings were more frequent in non-edge than edge contexts (11 vs. 5) and on unpaved than paved roads (11 vs. 5).

Stone martens (36%), European badgers (34%), and red foxes (30%) accounted for all dispersal events in scats. Additionally, carnivores comprised 44% of animal camera trap detections (359) and accounted for ∼70% of the 153 fig removals recorded, specifically by red foxes, stone martens, and Egyptian mongooses.

#### 3.2.1 Carnivore-mediated seed dispersal distances

We found no evidence that road-forest context or road type influenced ‘seed’ dispersal distances (Table 3; Fig. 4), contrary to our expectation of greater dispersal in non-edge contexts and along unpaved roads. Instead, distance from the road was positively associated with dispersal distance, with ‘seeds’ found 600 m from the road having traveled three times farther than those recovered just 5 m away.

**Figure 4.**
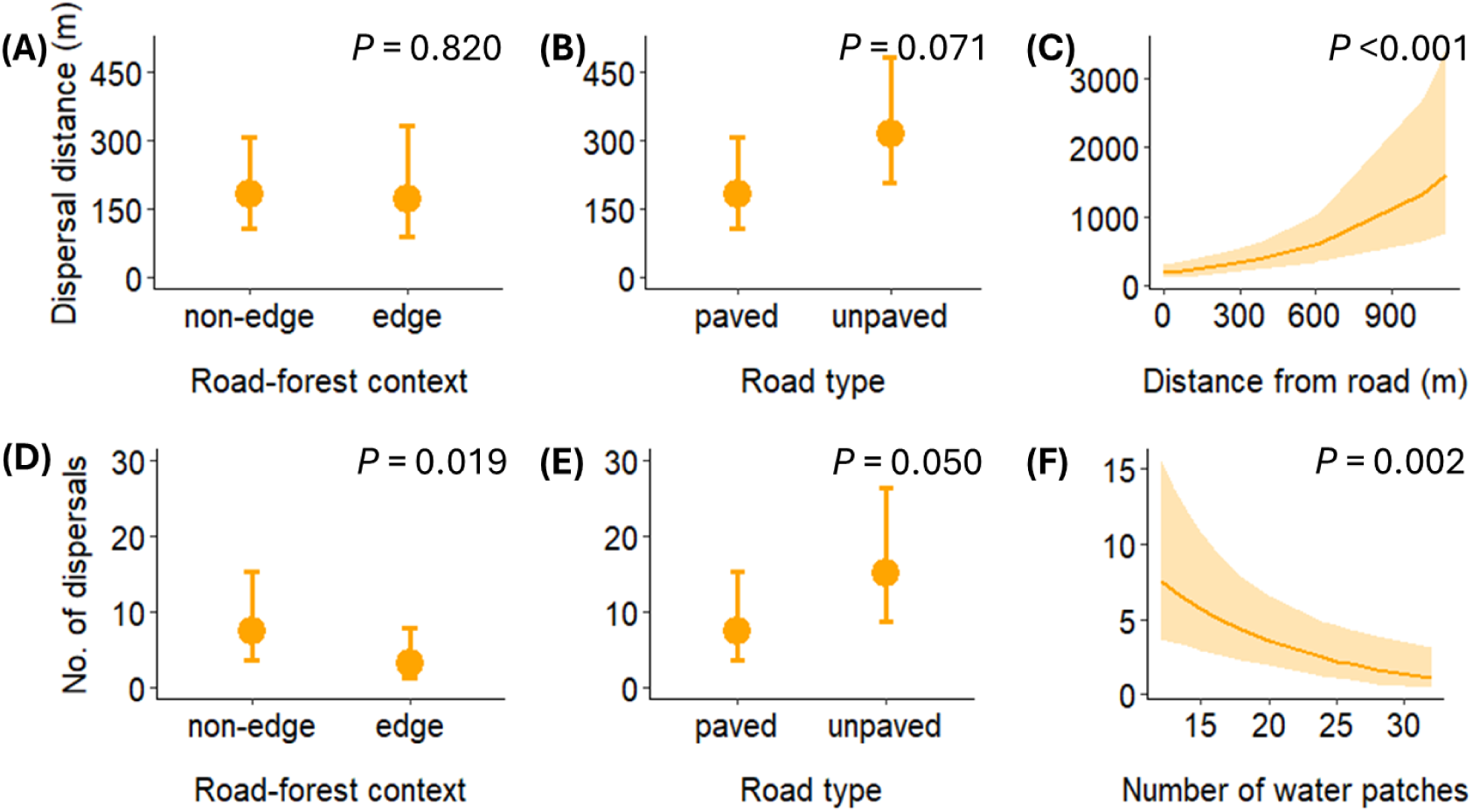
Mean fitted values (±95% CI) from the optimal mixed-effects models predicting (A–C) seed dispersal distance and (D–F) number of dispersals mediated by carnivores.

**Figure 5.**
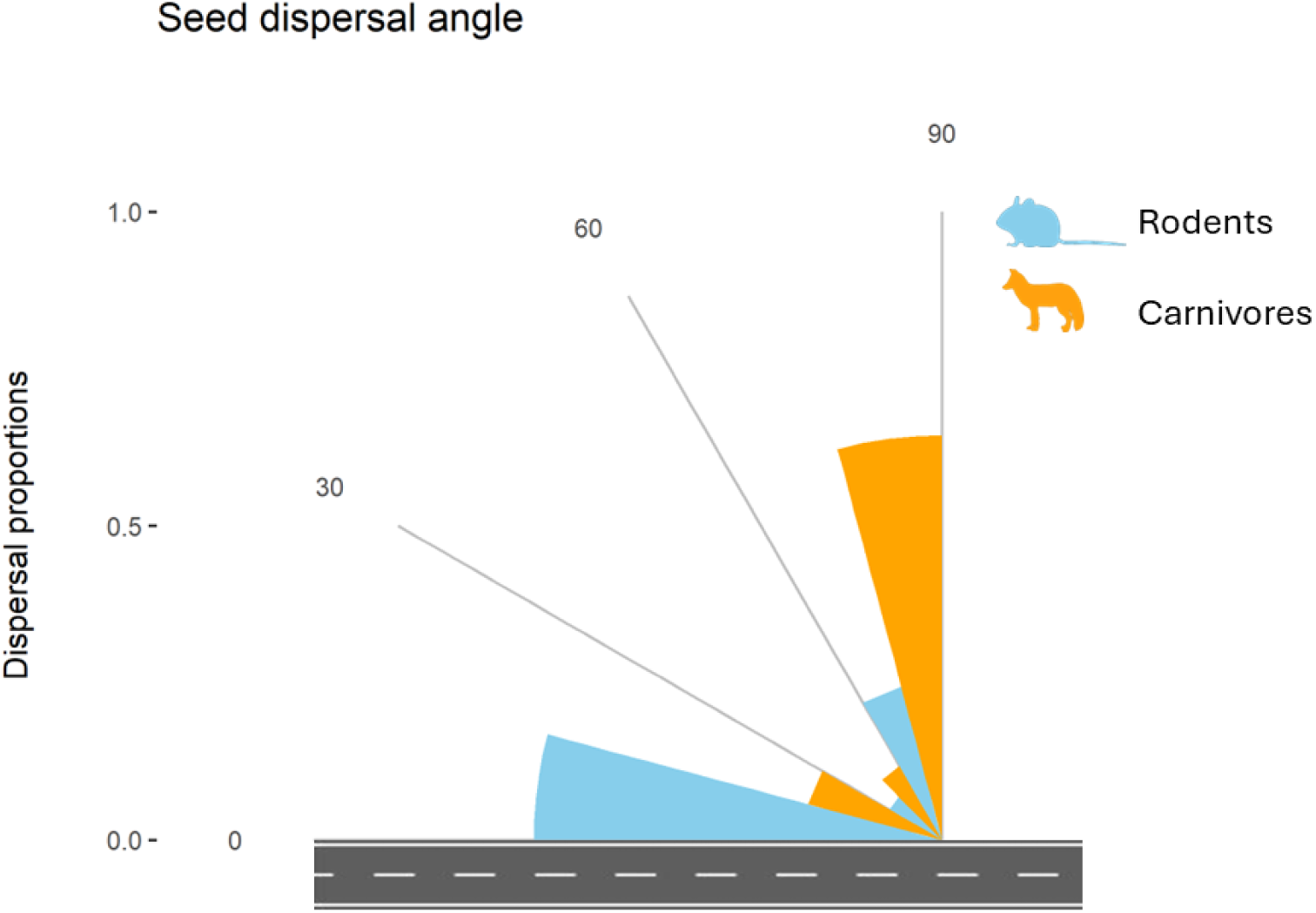
Directional patterns of seed dispersal by rodents (blue) and carnivores (orange) from supply stations located at roadsides.

#### 3.2.2 Number of carnivore-mediated seed dispersals

Consistent with our hypotheses, ‘seed’ dispersal events were more frequent in non-edge than edge road-forest contexts and at unpaved than paved road sites (Table 3, Fig. 4). Specifically, dispersal events were 2.3 times more numerous in non-edge contexts and twice as frequent at unpaved road sites. Additionally, the number of water patches within the 1200 m buffer significantly reduced dispersal frequency, with a 75% decline as water patches increased from 15 to 30.

### 3.3 Rodent- and carnivore-mediated seed dispersal direction

Most rodent-mediated acorn dispersals (65%) occurred along the road (0–30°), a proportion 1.9 times higher than the combined proportion at intermediate (30–60°; 10%) and perpendicular angles (60–90°; 25%), supporting our hypothesis that rodents would primarily disperse seeds along roadsides. In contrast, as predicted, carnivore-mediated ‘seed’ dispersals were predominantly perpendicular to the road (60–90°; 64%), occurring 1.8 times more often than at intermediate (30–60°; 14%) or along-road angles (0–30°; 22%).

## 4. DISCUSSION

The influence of road networks on seed dispersal in human modified landscapes remains poorly understood. By analyzing how two functionally distinct mammal groups—rodents and carnivores—mediate seed dispersal across different road-forest contexts, we show that roads influence not only dispersal distances and frequencies, but also dispersal directions and the likelihood of road crossings. While previous research has largely considered roads as barriers or corridors for wildlife movement (López-Barrera et al., 2007; Suárez-Esteban et al., 2013a; Chen et al., 2019a), our results reveal that road surface (paved vs. unpaved) and road-forest configuration (edge vs. non-edge) shape seed dispersal dynamics in complex ways. These findings provide a mechanistic understanding of how road network alters plant-animal interactions, with direct implications for landscape connectivity, ecological restoration, and road planning in Mediterranean oak forests.

Rodent-mediated dispersal was strongly influenced by road-forest context, with seeds moving farther at edge than non-edge sites, supporting our hypothesis that rodents disperse seeds over longer distances in edge habitats, where acorn availability is lower. Given that rodents commonly store acorns under shrubs (García-Hernández et al., 2016; Vaz et al., 2024), the lower shrub cover along road and roadless forest edges, especially on the open side, likely forces them to move seeds farther in search of suitable caching sites. Morán-López et al. (2016) suggested that in fragmented landscapes, rodents may prioritize acorn consumption over dispersal due to increased predation risk in open habitats. However, our results indicate that, despite potential risks, rodents still disperse seeds over longer distances in forest edges, suggesting that the benefits of securing acorns in protective microhabitats may outweigh the costs of increased movement. Studies have reported contrasting patterns of seed removal at edges, with some showing lower removal rates at sharp edges bordering pastures (López-Barrera et al., 2007), while others found higher removal rates in edge habitats, including by wood mouse (Mazzamuto et al., 2018). However, these studies did not account for the presence of roads. Our results suggest that roads themselves can amplify seed removal, highlighting the need for further research on how road-forest interactions shape seed fate across different landscapes.

For carnivores, road-forest context primarily influenced the number of dispersal events. Given their large home ranges and long gut-retention times (Jordano et al., 2007; Salgueiro et al., 2019; Rubalcava-Castillo et al., 2020), these animals readily disperse seeds across varied landscape patches, including open habitats and forests (Matías et al., 2010; Escribano-Avila et al., 2014; Rubalcava-Castillo et al., 2020). However, their lower activity in edge areas likely explains the reduced seed dispersal observed in our study. Species such as mongooses actively avoid open habitats (Palomares & Delibes, 1993), and pasture and cropland management in our study area likely further disrupts carnivore movement, thus limiting their role as seed dispersers. Research in central Germany has shown that seminatural habitats enhance the visitation and seed dispersal rates of species such as red foxes, stone martens, and badgers, whereas intensively used farmland leads to a decline in these rates (Grünewald et al., 2010). Understanding how different land uses influence carnivore movement is essential for designing landscape-scale connectivity strategies that sustain plant-animal interactions and maintain ecological functions and ecosystem services in fragmented forests (Phillips et al., 2020).

Road type played a key role in shaping seed dispersal patterns, with contrasting effects on rodents and carnivores. Unexpectedly, rodent-mediated dispersal distance was greater on paved than unpaved road sites, challenging our assumption that unpaved surfaces would better support small mammal movement. However, while Niu et al. (2021) reported longer dispersal distances near paved roads than within forest interiors, Chen et al. (2019a) found a 50% reduction in dispersal distance near paved roads, with this effect diminishing deeper into the forest. By directly comparing paved and unpaved roads, as well as roadless control sections, our study provides new insights into how road infrastructure influences rodent movement. Rodent densities were 1.6 times higher along both paved and unpaved roads than in roadless controls, suggesting that roads concentrate rodent activity within roadside verges. This is further supported by our finding that rodent seed dispersal occurred predominantly parallel to roads, a pattern not previously documented. This directional pattern of seed dispersal may help explain the high density of oak seedlings observed along roadsides in other studies (Woziwoda et al., 2018). The localized increase in rodent presence may explain the longer dispersal distances and greater seed movements among sparse shrubs observed near roads, particularly paved ones, reinforcing the role of roadsides as narrow but functionally significant seed dispersal corridors.

Despite higher rodent densities near roads, seed crossings remained rare, suggesting that roads restrict seed movement regardless of increased rodent activity. However, crossings were 2.1 times more frequent on unpaved than paved roads, indicating that paved roads impose greater movement constraints on small mammals (Ji et al., 2017; Grilo et al., 2018). This barrier effect is further highlighted by the sharp reduction in crossings on paved roads compared to roadless controls, whereas crossings on unpaved roads were three times more frequent than in their controls. These findings suggest that while roads concentrate rodent activity on roadsides, they still act as barriers to seed transport. However, road permeability varies with surface type, which has important implications for maintaining landscape connectivity in fragmented forests.

Carnivore-dispersed seeds crossed roads nine times more frequently than rodent-dispersed seeds, confirming roads as weaker barriers for carnivore-mediated dispersal. However, unpaved roads were particularly favorable, facilitating more seed crossings and increasing both dispersal frequency and distance. This supports the idea that unpaved roads function as seed dispersal corridors for larger mammals (Suárez-Esteban et al., 2013a; 2013b; Suzuki & Saito, 2023; Camacho et al., 2017), likely more so than paved roads. Carnivores preferentially disperse seeds along unpaved roads, likely due to reduced traffic and lower disturbance levels, which make these roads more suitable for movement and territorial marking (Suárez-Esteban et al., 2013b; Bueno et al., 2021). Future research should assess whether this pattern holds when comparing unpaved roads with traffic levels similar to those of paved roads. Indeed, paved roads can disrupt carnivore-mediated movements through direct mortality (Planillo et al., 2018; Silva et al., 2019), although some generalist carnivores, such as red foxes, may tolerate paved roads better than other species, particularly where roadside verges provide prey resources (Ruiz-Capillas et al., 2013; Galantinho et al., 2020).

Beyond road type and road-forest context, other factors also shaped seed dispersal. Acorn weight, arrival microhabitat, and shrub cover significantly influenced dispersal frequency and distance, aligning with previous studies (Gómez et al., 2008; Perea et al., 2011a; Zeng et al., 2022; Vaz et al., 2024). Rodents preferentially cache larger acorns (Cao et al., 2016), potentially increasing dispersal distances, though subsequent consumption often limits seedling establishment. Shrub cover was especially important, as acorns under shrubs dispersed farther than those in open habitats. Shrubs offer critical shelter and corridors for rodent movement along roadside verges, namely in Mediterranean oak forests (Rosalino et al., 2011; Galantinho et al., 2022), reinforcing their role in dispersal dynamics. Additionally, multiple water patches were associated with reduced carnivore-mediated seed dispersal, possibly due to riparian corridors diverting carnivore activity from roadsides (Ferreira et al., 2022; Santos et al., 2011; 2016). We also demonstrate for the first time that carnivores preferentially disperse seeds into forests, perpendicular to roads, rather than along roadsides.This highlights the need to manage roadside vegetation to support forest regeneration and connectivity. Indeed roadsides may serve as seed sources rather than just movement corridors, influencing regeneration patterns in adjacent forests.

Considering these findings, enhancing mammal-mediated seed dispersal in human-modified landscapes requires targeted roadside vegetation management, particularly in edge habitats, which already border 70% of the world’s forests (Bhatt et al., 2023). Roadside verges are increasingly recognized as valuable conservation areas, providing habitat, movement corridors, and even nurse sites for plant regeneration (Woziwoda et al., 2018). However, their management must balance ecological benefits with potential trade-offs, such as creating ecological traps or facilitating invasive species, which can undermine conservation goals (Mazzamuto et al., 2018; Phillips et al., 2019). To maximize their ecological functionality, management strategies should account for the differential effects of road surface types on mammal dispersers. For rodents, priority should be given to roadsides on paved roads, ensuring moderate shrub cover to provide rodents with safe caching sites and reduce predation risk, while preventing excessive shrub encroachment through periodic mowing. This approach can facilitate rodent-mediated seed dispersal and promote forest recovery without forgetting that unpaved roads promote more acorn crossings, likely due to lower traffic volumes and disturbance levels, thus offering greater potential for forest expansion in edge contexts than paved roads. For carnivores, unpaved roadside verges should be prioritized, with management efforts focused on creating vegetation corridors that connect forests fragmented by open landscapes. By tailoring interventions to the distinct ecological roles of rodents and carnivores, road networks can be managed not only to reduce their barrier effects but also to actively support landscape connectivity and drive forest regeneration in Mediterranean oak ecosystems.

## 5. CONCLUSIONS

Our study demonstrates that roads influence seed dispersal dynamics in Mediterranean oak woodlands through the combined effects of road type, road-forest context, and the ecological roles of rodents and carnivores. Rodents, while primarily dispersing seeds locally—often along roadsides where they were more abundant—dispersed them farther in edge forest contexts and near roads, particularly paved ones, possibly influenced by acorn availability and shrub cover. In contrast, carnivores facilitated long-distance dispersal, predominantly perpendicular to roads, and were more likely to cross roads, especially unpaved ones, though their frequency of seed dispersals was influenced by landscape features such as water patches. Importantly, we show that roads are less of a barrier to carnivore-mediated seed dispersal than to rodent-mediated dispersal. These findings reveal that roadsides act not only as movement corridors but also as potential seed sources, with implications for forest regeneration and landscape connectivity.

These insights underscore the importance of integrating ecological processes into road planning and management. For rodents, maintaining moderate shrub cover along paved roadsides can enhance seed dispersal and promote forest recovery. For carnivores, prioritizing unpaved roads and creating vegetation corridors can support long-distance seed movement and connectivity across fragmented landscapes. By tailoring management strategies to the distinct roles of these dispersers, road networks can be designed to reduce their ecological impacts and actively contribute to the ecological functionality of Mediterranean oak ecosystems. Recognizing the complementary roles of rodents and carnivores in seed dispersal further highlights the potential of road networks to support both local and long-distance dispersal, enhancing the resilience of these ecosystems.

## CRediT authorship contribution statement

**João Craveiro**: Data curation, Formal analysis, Investigation, Visualization, Writing – original draft. **Miguel N. Bugalho**: Supervision, Writing – review & editing. **Pedro G. Vaz**: Conceptualization, Supervision, Resources, Writing – review & editing.

## Acknowledgments

JC was supported by the Portuguese Science and Technology Foundation (FCT) through an individual grant (2021.05551.BD) and institutional funding to CEABN-InBIO (DOI 10.54499/UIDB/50027/2020) and InBIO (DOI 10.54499/LA/P/0048/2020). JC and PGV were funded by FCT through CE3C (DOI 10.54499/UIDB/00329/2020) and CHANGE (DOI 10.54499/LA/P/0121/2020). PGV received funding from the AdaptForGrazing project (PRR-C05-i03-I-000035-LA4.3/4.4/4.6/4.7), supported by the EU (RRF) via IFAP (PRR). We thank Simone Erroi and Abdullah Ibne Wadud for their assistance during pilot studies.

